# Viral lifecycle dynamics and spatial structure explain why accessory genes are associated with temperate phages

**DOI:** 10.1101/2025.02.28.640786

**Authors:** Sultan A. Nazir, Bram van Dijk

## Abstract

Genes encoding for virulence factors are frequently found on prophages, yet the evolutionary forces driving this association remain unclear. The evolutionary association of mobile genetic elements (MGEs) with host-beneficial genes is known to be hindered by the stability of chromosomal loci and competition with MGEs lacking accessory genes. Using a mathematical model that incorporates these constraints, we identify two key mechanisms that help overcome them, resulting in the evolutionary linkage of accessory genes, like virulence genes, to prophages. First, we show that migration across bacterial populations favours the association of genes with prophages when the gene is beneficial in certain environments (e.g. virulence genes in the gut) that also trigger higher rates of prophage induction. Second, we show that within-population spatial dynamics also promotes the association of weakly selected genes and phages. Here, virion dispersal allows phage-encoded genes to spread into patches of bacteria lacking the gene, giving them a selective advantage over immobile chromosomal genes. We argue that these mechanisms are less applicable for plasmids and other MGEs, highlighting a potentially unique role for phages in shaping bacterial adaptation. By demonstrating how phage lifecycle dynamics and spatial heterogeneity drive MGE-gene associations, our work provides new insights into the evolution of phage-encoded virulence.

## Background

Mobile genetic elements (MGEs) are genome segments that can move within or between genomes, often replicating independently of the chromosome. Although MGEs such as bacteriophages, plasmids, and transposons have been the focus of extensive research for several decades, their eco-evolutionary functions are still being elucidated (Mark Osborn and Böltner 2002; Hall, Harrison, and Baltrus 2021). Increasingly, studies reveal the crucial roles that MGEs play in shaping the ecological and clinical functions of microbial communities (D. J. Rankin, E. P. C. Rocha, and S. P. Brown 2011; Weisberg and Chang 2023). MGEs are especially relevant because they can carry accessory genes, which are non-essential for the MGE itself, but may substantially alter the bacterial phenotype. Different types of MGEs appear to vary in their likelihood of carrying specific accessory genes (D. J. Rankin, E. P. C. Rocha, and S. P. Brown 2011), but the mechanisms behind these associations remain poorly understood. For example, a recent analysis shows that phages are more likely to encode virulence factors than plasmids (Takeuchi, Hamada-Zhu, and Suzuki 2023). This raises the question: what makes phages uniquely predisposed to turn their hosts into pathogens? Understanding the conditions under which virulence genes become phage-borne is critical, as this enables horizontal transfer to other bacterial strains and potential disease outbreaks (She et al. 2025; Davies et al. 2015).

Several hypotheses have been advanced to explain why non-essential or cooperative traits are frequently found on MGEs rather than on chromosomes (D. J. Rankin, E. P. C. Rocha, and S. P. Brown 2011). For phage-encoded pathogenicity specifically, Abedon and LeJeune provide an excellent review of proposed and empirically supported mechanisms (Abedon and LeJeune 2007). A leading hypothesis emphasizes epistasis through molecular coregulation of phage and virulence genes. Indeed, in many systems these genes are tightly coregulated, either by shared transcription factors, such as csrA in S. enterica (She et al. 2025), or through prophage induction serving as a trigger for exotoxin release, as in the case of Shiga toxin in E. coli (Rodríguez-Rubio et al. 2021). However, phage-borne virulence genes are not universally coregulated with prophage induction, and several studies report independent regulation from lysis genes, where induction triggers do not affect virulence expression (Brouwer et al. 2020; Banks, Lei, and Musser 2003). Thus, while coregulation can be sufficient to explain phage–virulence linkage, it is not necessary.

Models have been a useful tool to unpack the conditions under which MGEs carry certain accessory genes. For example, Bergstrom, Lipsitch, and Levin proposed a local adaptation hypothesis, wherein plasmid-encoded genes persist by transferring locally adapted genes to newly arriving strains (Bergstrom, Lipsitch, and Levin 2000). Later, Smith proposed the infectivity hypothesis, which stated that MGEs may evolve to carry genes for public goods in order to force defectors to cooperate (Smith 2001). More recently, van Dijk et al. have shown that horizontal gene transfer (HGT) can rescue a beneficial gene (therewith increasing population fitness) when the benefit is not strong enough to maintain the gene against mutational loss (van Dijk et al. 2020). From these studies, we can generalise a *rescue hypothesis* (Dijk 2020): genes are MGE-encoded when HGT can rescue the gene from being lost through negative selection, mutational loss, or the emergence of cheaters.

While the above-mentioned models and the rescue hypothesis are insightful, they do not fully capture the biology of how MGE-driven mobilisation comes about. For an accessory gene to become mobilised by residing on an MGE, its mobility must not only be beneficial, it must also persist under competition against MGEs that do not encode the accessory gene. Competition with such “non-carrier” MGEs has indeed been shown to hinder the association of MGEs with accessory genes (Geoffroy and Uecker 2023; Mc Ginty, Daniel J. Rankin, and Sam P. Brown 2011). In addition, most mechanisms of gene loss will act on the mobile locus as much as they do on the chromosomal locus, and the gene may fail to persist by mobility alone.

More recent work by Lehtinen, Huisman, and Bonhoeffer has shown that an accessory gene that is initially more abundant on the mobile (MGE) locus than on the immobile (chromosomal or a cryptic prophage) locus will, under weak selection, stay on the mobile locus (Lehtinen, Huisman, and Bonhoeffer 2021). This “priority effect” may emerge when a species is more likely to acquire the gene via an MGE than by mutational innovation on the chromosome. While this mechanism is likely of importance to plasmids which have host ranges that span broad taxa (Coluzzi and Eduardo PC Rocha 2025), temperate phages are generally thought to have more narrow host ranges, often limited to strains of a single species, due to their specificity during integration (Bobay, Eduardo P.C. Rocha, and Touchon 2013; Bobay and Ochman 2018; Moura de Sousa et al. 2021). Moreover, there is evidence that genes associated with prophages have a chromosomal origin due to their localisation close to the attachment site of the prophage DNA (Brüssow, Canchaya, and Hardt 2004) and based on phylogenetic analysis (She et al. 2025). In other words, while much research on MGEs was done with plasmids as a model MGE, implicitly assuming generalisability to other MGEs, fewer studies have explicitly modelled other vectors of HGT.

In this article, we focus on understanding why a beneficial chromosomal gene can evolve to be associated with bacteriophages by means of mathematical modelling. While our model could be applied to accessory genes more generally, we built our model especially in the context of exotoxin production, a very common virulence factor encoded by prophages (Boyd and Brüssow 2002; Gummalla et al. 2023). The benefits provided by these extracellular virulence factors may be partially enjoyed by other bacteria that also gain access to resources of the plant or animal through tissue damage. Therefore, we model the expression of the gene as a cooperative trait with a certain degree of privatisation of benefits in order to maintain a positive selection pressure. As a preliminary analysis, our mathematical model verifies that the “rescue” of certain beneficial genes by lysogenic phages is highly sensitive to competition with non-carrier phages. In competition with non-carrier phages, we find that phage-encoded genes cannot invade when the gene is already chromosomally encoded. However, we find that the gene-phage association can invade under certain conditions within two distinct implementations of spatial structure. Firstly, by extending our Ordinary Differential Equation (ODE) model to include a host-environment metapopulation structure, we find that when prophage induction is upregulated in the host environment (where the benefits of the gene are limited to), natural selection favours the beneficial gene to be phage-encoded. Secondly, using an individual-based model, we propose virion dispersal within a spatially structured population as a mechanism that catalyses the evolution of phage-encoded cargo genes under weak selection. Here, since natural selection occurs at a local scale, phage particles help the gene disperse to pockets lacking the gene, reinforce selection, and replace the chromosomal gene at a global scale. Taken together, our work shines light on the *exogenous* factors (in contrast to endogenous factors such as molecular coregulation) that contribute to the evolution of phage-encoded accessory traits, explaining the observed association of virulence genes and prophages.

## Methods

### Infection cycle and genotype space

We implemented an infection model, as shown in Figure 1(A), where uninfected prokaryotic cells (denoted by *U*) are susceptible to infection by virion particles (denoted by *V*) such that the viral DNA integrates into the cell as a prophage and converts each cell into a lysogen (denoted by *L*). The prophage is then induced at a rate *α* which causes cell lysis and the release of virion particles equal to its burst size *r*_*V*_. Assuming that they occur simultaneously, we ignore the time lag that occurs between infection and integration and between induction and lysis. Virion particles in the external environment can either decay at a rate *δ* or infect a susceptible cell at a rate *β*. In this framework, we assume superinfection immunity such that a cell cannot be infected by a virion particle if it already harbours a prophage.

**Figure 1.**
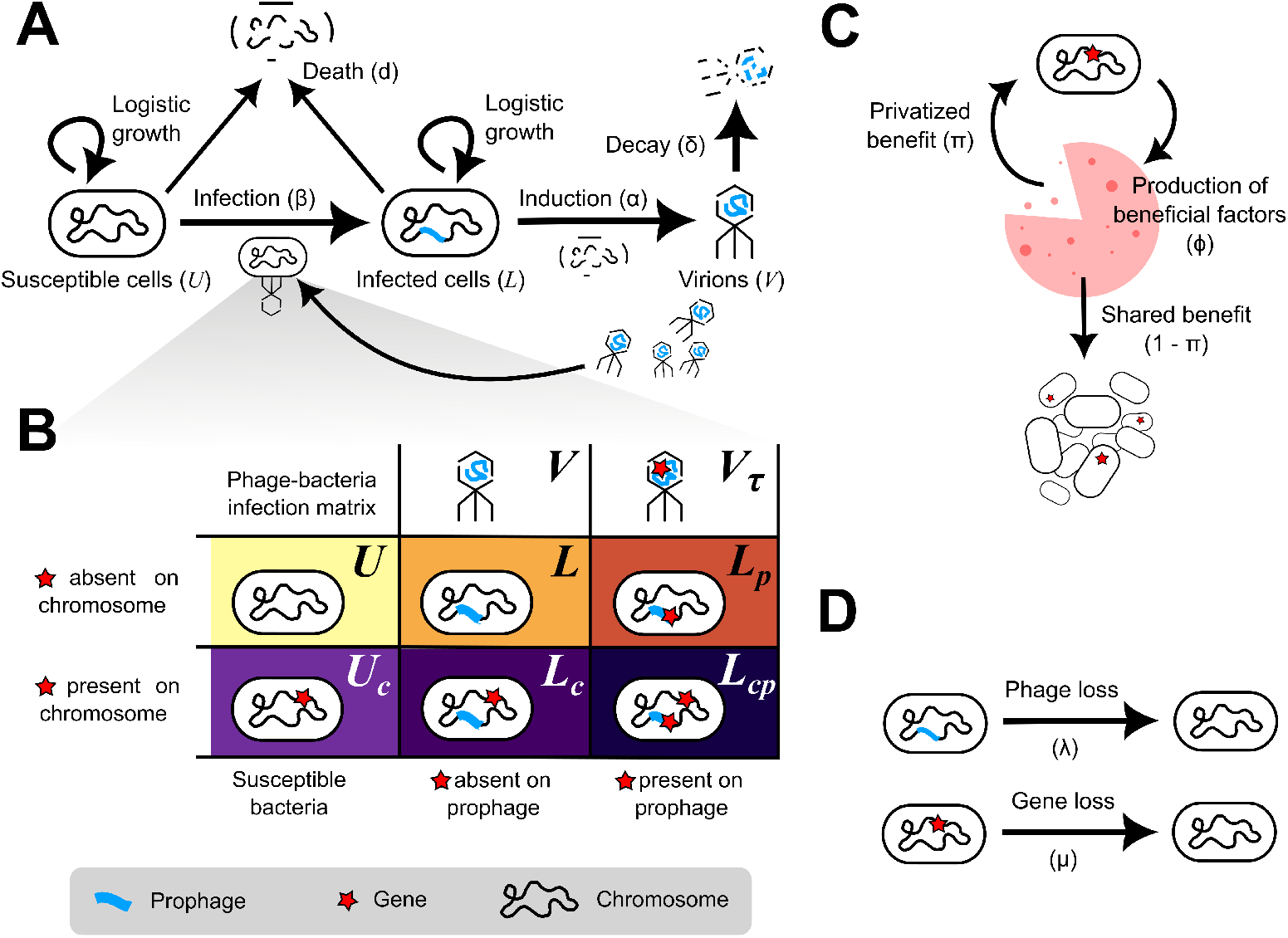
An illustration of all of the model components: (A) The lifecycle of the phage consists of a prophage stage that begins upon infecting a susceptible bacterium and a virion state which is produced upon induction of the prophage. As a virion, the phage can be lost to environmental degradation. (B) Based on whether the infecting phage carries the gene of interest or not, and whether the susceptiple cell carries the gene at the chromosomal locus, we get six cell genotypes and four of them carry the gene. The subscript denotes whether the gene is present on the chromosomal locus (*c*) or the phage locus (*p*). (C) Out of the *ϕ* benefits produced by a carrier, *π* fraction is privatised by the producer and the remaining is shared with the whole cell population. (D) Both the gene and the phage can be lost from a microbial genome due to mutations.

The viral DNA may or may not carry the gene of interest. If the virion particle is a carrier of the gene, it is marked by a subscript *τ*. When a non-carrier virion *V* infects a susceptible cell, it creates a non-carrier prophage, but if a carrier virion *V*_*τ*_ does, it creates a carrier prophage, and a lysogen with a carrier prophage is indicated by a subscript *p* to denote that the gene is encoded by the phage. Every cell, regardless of whether it is infected by a prophage or not, has a chromosomal locus where the gene may or may not be present. If the cell is a carrier of the gene at the chromosomal locus, it is indicated by a subscript *c*. Hence, as shown in Figure 1(B), there are six possible genotypes of prokaryotic cells (*U, U*_*c*_, *L, L*_*c*_, *L*_*p*_, and *L*_*cp*_) and two kinds of virion particles (*V* and *V*_*τ*_).

### Cooperation and privatisation

All cells grow at a fixed rate and compete for resources along with an intrinsic death rate of *d*. The effect of carrying the gene of interest on the growth dynamics is modelled as follows. We define “minimum carrying capacity” as the carrying capacity in the absence of the gene and set its value to one for non-dimensionalisation, i.e. all population sizes are expressed relative to the minimum carrying capacity. Let *ϕ* represent the maximum additional biomass that can be generated due to gene expression. If *f*_*τ*_ denotes the fraction of carriers in the population, by the definition of *ϕ*, the net carrying capacity becomes 1 + *ϕf*_*τ*_. If the benefits produced by the carriers are equally shared among all cells, each cell receives a benefit of *ϕf*_*τ*_. However, positive assortment and other mechanisms enable cooperators to have disproportionate access to these benefits. To model this, as shown in Figure 1(C), we introduce the parameter *π*, the fraction of the benefits produced by a carrier that is privatised, i.e. retained exclusively by the producer. Consequently, all cells (including the carriers themselves) receive a shared benefit of (1 − *π*)*ϕf*_*τ*_, while the carriers gain an additional private benefit of *πϕ*. This framework allows *π* to function as a “privatisation parameter”, or more generally, to tune the rate of selection acting on the gene.

### Mutations

In the model, we consider two kinds of deletion mutations, as shown in Figure 1(D). Firstly, every “carrier” (henceforth used to refer to a cell that has the gene of interest) can mutate into a “non-carrier” by gene loss at a rate *µ*. Note that this mutation occurs at the prophage locus and the chromosomal locus, at the same rate. Secondly, every lysogen can lose its prophage at a rate *λ*. For simplicity, we assume that this loss coincides with the loss of superimmunity, i.e. the cell will immediately be available for future phage infections. If a cargo gene resided on the prophage, this gene is lost along with the phage.

The above processes are first analysed within a framework of differential equations (Supplementary Information A), where we focus on the ability of a phage-encoded trait to replace a chromosomally-encoded trait. In addition, this analytical model is compared and contrasted with an individual-based framework, described in the next section. The parameters and their values are summarised in Supplementary Information B.

### Spatial model

For the spatially structured version of the model, we implemented an Individual based model (IBM) of a finite population of bacteria on a square grid. The model captures the same processes described above using the ODEs while introducing spatial structure and finiteness to the cell and virus populations with the following elements.

### Birth and Death

A cell can copy itself within a neighbouring empty site with a probability *b* as shown in Figure 2(A). We assume that the rate of birth is equal for all cells at all times. The effect of cooperative interactions is expressed in terms of varying death rates. We chose to vary the death rates instead of the birth rates, because birth is inherently density-dependent due to our assumption that reproduction requires empty sites. In the absence of any benefit from carriers, a cell dies with probability *d*_*s*_ + Δ*d* where *d*_*s*_ is the intrinsic death rate in the spatial model and Δ*d* is the rate of death that can be avoided with the benefits from the gene. The maximum population size is limited by the size of the grid (total number of grid points). Therefore, to simulate dynamics comparable to the ODE, we adjust the death rate parameters such that the grid size (the maximum carrying capacity) is 1 + *ϕ* times the average steady population size in the absence of the gene (the minimum carrying capacity). Analogously to the ODE model, the benefits of the cooperative gene are partially privatised. A carrier privatises *π* fraction of its benefits to itself (lowering its own chance of death), with the remaining fraction (1 − *π*) of benefits shared with others. Therefore, the probability of death is

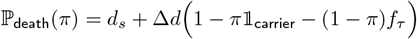

where 𝟙_carrier_ is the indicator variable

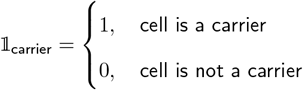

and *f*_*τ*_ is the fraction of carriers on the grid.

**Figure 2.**
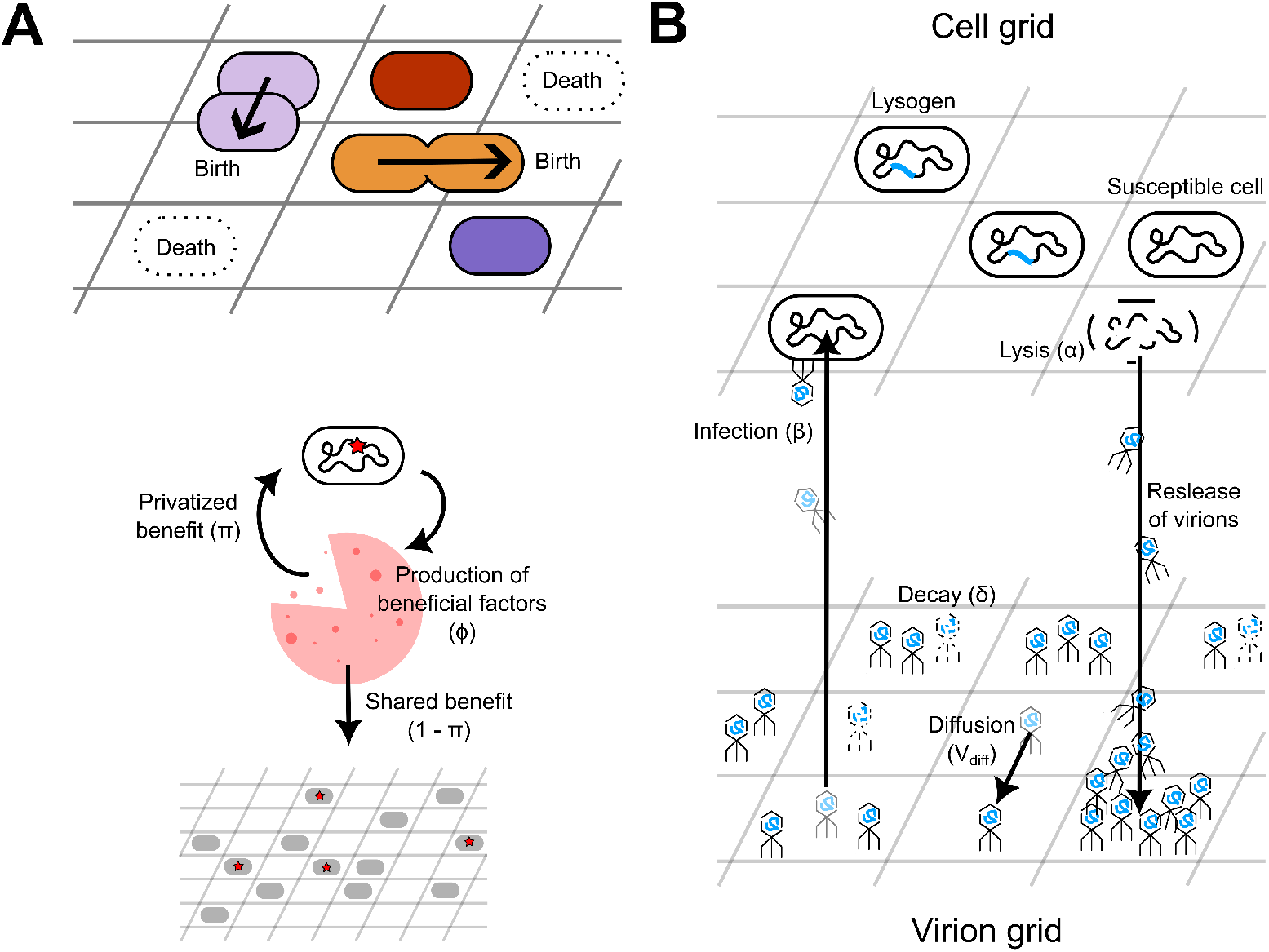
Spatial model: (A) Individual cells are placed on a square grid and the occupied grid points are coloured according to the cell genotype (identical to the colours in Figure 1(B). On the grid, the cells can reproduce into an unoccupied grid point in its Moore neighbourhood. The cells can also die, leaving an unoccupied grid point. The propensity of death reduces with the beneifts received from the gene. If a cell is a carrier, it privatises *π* fraction of benefits and shares the remaining 1− *π* fraction with other cells on the grid. (B) Apart from the ‘Cell grid’, there is a ‘Virion grid’ where more than one virion particle can occupy a single grid point. They infect cells and undergo part of their lifecycle as prophages on the Cell grid. Upon induction and lysis, virions are released onto the Virion grid where it can diffuse and ultilmately infect a susceptible cell or undergo decay.

To ensure that the ODE is a good approximation of the spatial model when well-mixed, all rate parameters in the IBM are scaled by a factor of 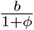 (for example, 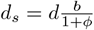) and we set 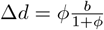. See Supplementary Information C for verification of the well-mixed approximation.

### Phage lifecycle

Extracellular virions are also modelled as discrete entities. Therefore, at every grid point, we have an integer measure of the number of extracellular virion particles of each type – non-carrier and carrier. Upon induction and lysis, the cell releases the virion particles into the ‘Virion grid’ at the same grid coordinate as shown in Figure 2(B). The released virions are carriers if the induced prophage was a carrier. Irrespective of whether it is a carrier or not, at every time point, each virion particle can decay or migrate to a neighbouring grid point with fixed probabilities for each. The probability of migration *V*_diff_ controls the rate of diffusion of these particles. When a susceptible cell is located at a specific coordinate on the ‘Cell grid’, all virion particles present at the same coordinate on the Virion grid compete to infect the cell. Each virion particle has a fixed probability of infecting the cell. Therefore, the probability that the cell becomes infected increases linearly with the local virion abundance.

## Results

### Gene association with prophage does not evolve in a well-mixed population

To investigate eco-evolutionary outcomes where phages carry the beneficial gene as cargo from a chromosomal origin, we performed numerical simulations across the *π* − *α* parameter space. Since our model does not include gene translocation between the chromosome and the prophage, we simulate an invasion attempt of a mutant that encodes the gene on a prophage within a resident population where the gene is chromosomal, by setting the initial state to *L*_*c*_ = 1.98, *L*_*p*_ = 0.02, and the remaining genotype abundances to zero. In other words: all cells have a prophage, 99% of all cells encode the gene chromosomally, and the remaining 1% of cells have the gene encoded on a prophage instead. Hence, the question becomes whether the rare mutant encoding the gene on a prophage invades the population. There is a persistence region (see Figure 3(A)) — where both the phage and the gene can coexist. This is because the gene cannot persist if selection is not strong enough against gene loss, and the phage cannot persist if induction rate is not strong enough against phage loss (See Supplementary Information D for the derivations). Therefore, within the persistence region, we examined the association between the gene and the phage in terms of the percentage of prophages that carry the gene and the percentage of gene copies that occur on the prophage (shown with a 2D colour gradient).

**Figure 3.**
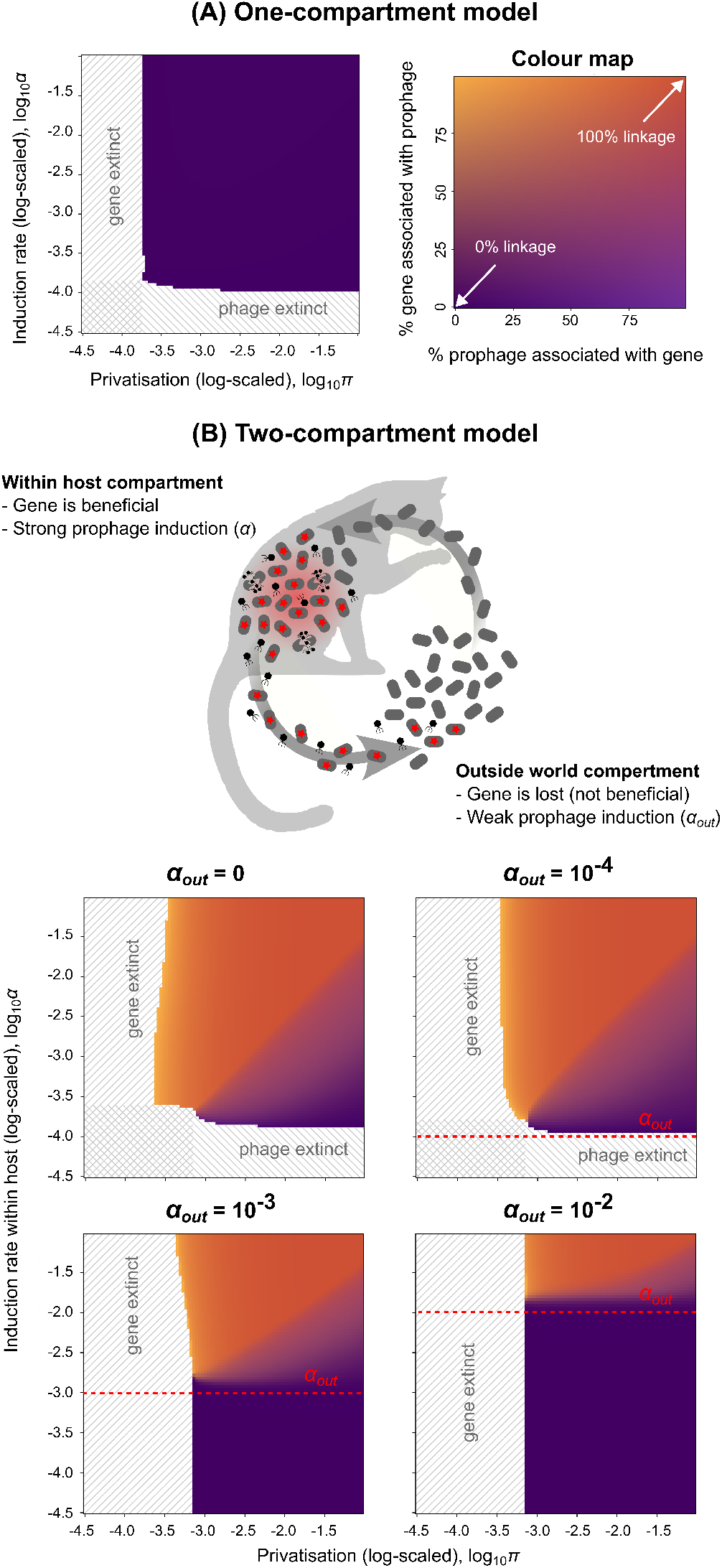
ODE model: (A) In an unstructured environment, i.e. one well-mixed compartment of cells, gene-phage association does not evolve. The degree of association of gene and phage (represented by the 2D colour map on the right) is plotted after simulation time 10^6^ for different values of *α* and *π*. (B) When a new compartment simulating the “outside world”, where the gene is not beneficial, is introduced, gene-phage association evolves but it requires that *α > α*_out_.

We observed that these conditions are not sufficient for the gene-phage association, since the mutant does not establish itself in the population (Figure 3(A) remains fully dark purple). This result indicates that, assuming genes with chromosomal origins, it is difficult for the gene to be genetically linked to phages as cargo. Hence, in agreement with previous models (Mc Ginty, Daniel J. Rankin, and Sam P. Brown 2011; Lehtinen, Huisman, and Bonhoeffer 2021), prophage-encoded (more generally MGE-encoded) accessory genes that are typically traced back to a chromosomal origin cannot evolve in a well-mixed population.

Why is it so difficult for a beneficial gene to become stably associated with a prophage? When the gene is initially located on the bacterial chromosome, infection by phages carrying the gene does not confer an additional fitness advantage to already gene-bearing cells, and thus these phages are not selectively favoured. Although susceptible cells lacking the gene can gain a fitness benefit upon infection by carrier phages, the overall propagation of the gene is limited by the presence of a competing population of non-carrier phages which are always present due to gene loss mutations. Additionally, the gene can be lost from the prophage due to phage loss mutations. As a result, the net growth rate of the phage-associated gene cannot surpass that of the chromosomally encoded gene.

### 1. Spatial structure between populations: differential induction promotes gene-prophage association

In their review, Abedon and LeJeune have explored endogenous factors such as epistasis that promote the association of beneficial genes with prophages (Abedon and LeJeune 2007). However, beyond within-cell biochemical interactions, the co-regulation of prophage induction and gene expression can also emerge from *exogenous* factors such as environmental heterogeneity. The gene may not be advantageous in all environments — for example, a pathogenicity trait is only beneficial within a host environment. Similarly, prophage induction is often influenced by extracellular conditions (Huisman, Bernhard, and Igler 2025). Henrot and Petit identified multiple signalling molecules within the mammalian gut microbiota that can trigger prophage induction (Henrot and Petit 2022). In addition, expression of benefits from the gene itself may alter the cellular environment in ways that promote induction. For example, Pattenden, Eagles, and Wahl propose that virulent bacterial strains, due to their faster growth and increased exposure to stress, induce their prophages more frequently (Pattenden, Eagles, and Wahl 2021). Furthermore, immune responses targeting virulent cells can inadvertently trigger prophage activation.

Motivated by these observations, we next consider a scenario in which the gene is beneficial only in a specific environment, and this same environment promotes prophage induction. To model this, we introduce a second compartment representing the “outside world” where toxin production does not confer a fitness benefit and prophage induction signals are weak (see Supplementary Information E for modified equations). In the absence of migration, the system in this outside world will converge towards a population of non-carrier susceptible (*U*) cells. We connect this compartment with the “host environment” by introducing migration between them. We assume that the migration is symmetric and that both compartments have equal maximum carrying capacities.

With this modification, we see a large portion of the parameter space permitting the invasion of phage-borne genes within the host environment (bright red colouration in Figure 3(B)). This is because the cells in the outside world are inclined to lose both the gene and the phage, and the influx of these non-carrier susceptible cells push the threshold of selection required to maintain the gene. This can be seen by the expansion of the ‘gene extinction’ region in the parameter sweep. Coincidentally, this influx increases the rate of phage-infections that successfully convert non-carriers into carriers within the host environment, hence rescuing the gene from extinction close to the threshold. If this rate of conversion by phage transmission can exceed the rate of selection replacing incoming non-carriers with carriers, there is a selection pressure for mobility at the gene-level. Hence, the bright red region appears for weak selection and a high rate of prophage induction. It is interesting to note that the gene-phage association is not a stable attractor when either of the compartments is considered to be a closed system. However, upon permitting migration between them, phage-association becomes the stable evolutionary attractor.

To highlight the requirement of a weaker rate of prophage induction in the outside world (*α*_out_), we repeated the simulations for different values of *α*_out_ (shown in red dashed line). We saw that the gene-phage association evolves only when the rate of prophage induction in the host environment exceeds *α*_out_. Therefore, the higher proportion of uninfected cells in incoming cells, due to the weaker transmission rate of the phage in the outside world, is a requirement for the emergent selection pressure on the mobility of the gene within the host environment.

More generally, we postulate the following hypothesis from our metapopulation framework.

*If patches where a gene provides higher fitness benefits also have higher mobile element transmission rates, this positive correlation creates a selection pressure for linkage of the gene and the mobile element*.

The analysis so far has been performed using a set of 16 differential equations (8 equations for each compartment), which assumes that the population is homogeneous within each compartment. However, natural microbial populations are also spatially structured *within* these patches, exhibiting local variations in bacterial density and gene/phage distribution. In the next section, we explore whether spatial structure within a patch alone can drive gene-phage associations without requiring additional regulatory mechanisms.

### 2. Spatial structure within a population: virion dispersal promotes gene-prophage association

In the individual-based spatial model (described in Methods), due to discrete population sizes, a gene can be lost entirely and not re-emerge. To investigate whether phage-encoded cargo genes can establish, we conducted an invasion experiment. We initialised a population of *L*_*c*_ cells on a 120 *×* 120 grid and allowed mutant types (*U, U*_*c*_, and *L*) to arise and stabilise their abundances. We then introduced a spontaneous translocation mutation, converting 1% of *L*_*c*_ cells into *L*_*p*_ cells, effectively moving the gene from the chromosome to the prophage. If the phage-associated gene went extinct, we reintroduced the mutation in the same manner. The simulation ran until either the gene was lost from both loci or the maximum time limit (5 *×* 10^6^ steps) was reached. A “successful invasion” was defined as the complete replacement of the chromosomal gene by the phage-associated gene.

Running 20 replicates of the experiment across different values of privatisation (*π*) and virion diffusion rates (*V*_diff_) revealed two key findings (Figure 4). First, phage-associated genes successfully invaded in some replicates under intermediate privatisation. While the ODE model never allowed full takeover of the phage locus, spatial simulations frequently did. Control experiments with well-mixed populations confirmed that this effect was significantly driven by spatial structure rather than finite population size or reintroduction of mutants (see Figure 4(B)). Second, increasing virion dispersal noticeably enhanced phage-associated gene invasion. However, when both cells and virions were well-mixed, virion diffusion no longer influenced gene-phage association, highlighting the role of differential dispersal in moving the phage-associated gene away from local competition with the chromosomal gene.

**Figure 4.**
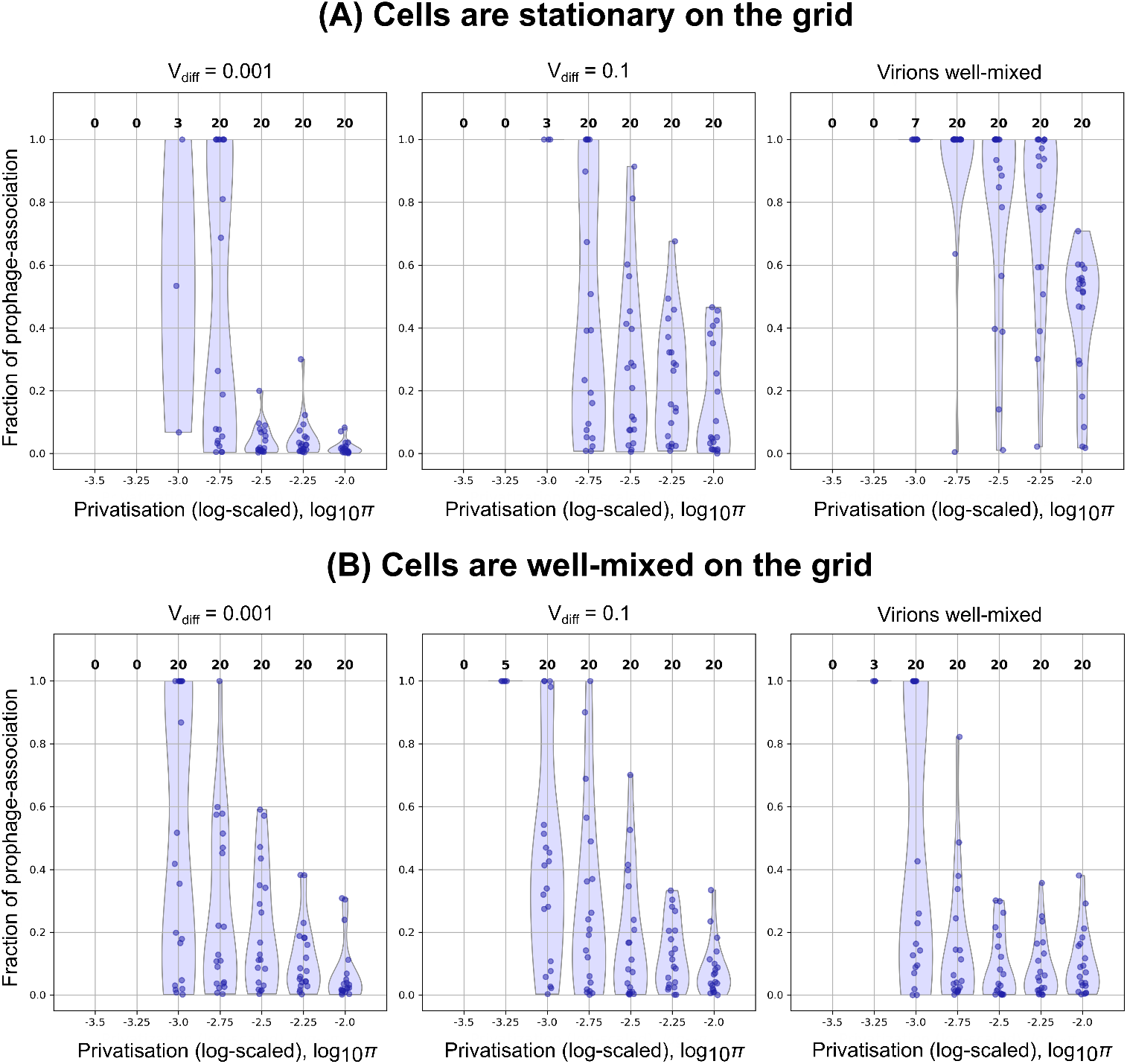
Differential dispersal of virions in a spatially structured cell population leads to the evolution of gene-phage association for intermediate selection. The plots show the distribution of final fraction of prophage-association in the gene population, excluding runs where the gene went extinct. The number above each strip-plot indicates the number of replicates where the gene survived. For each value of *π* and *V*_diff_, the distribution is plotted over 20 replicates of the stochastic simulation. (A) Cells are stationary on the grid. (B) Cells are well-mixed, i.e. the grid points on the ‘Cell grid’ are shuffled every time-step.

To better understand the conditions under which the phage-encoded gene can invade, we analysed spatial snapshots and temporal dynamics across different parameter regimes (Figure 5). In spatially structured environments, local reproduction leads to clustering of genotypes, forming spatial niches where the success of an invading genotype depends on local context. Carrier genotypes have a higher growth rate and are able to invade patches occupied by non-carrier cells (visible in orange and yellow in panels a and c). However, gene loss mutations continually regenerate non-carrier genotypes, allowing gene-free clusters to persist under weak selection.

**Figure 5.**
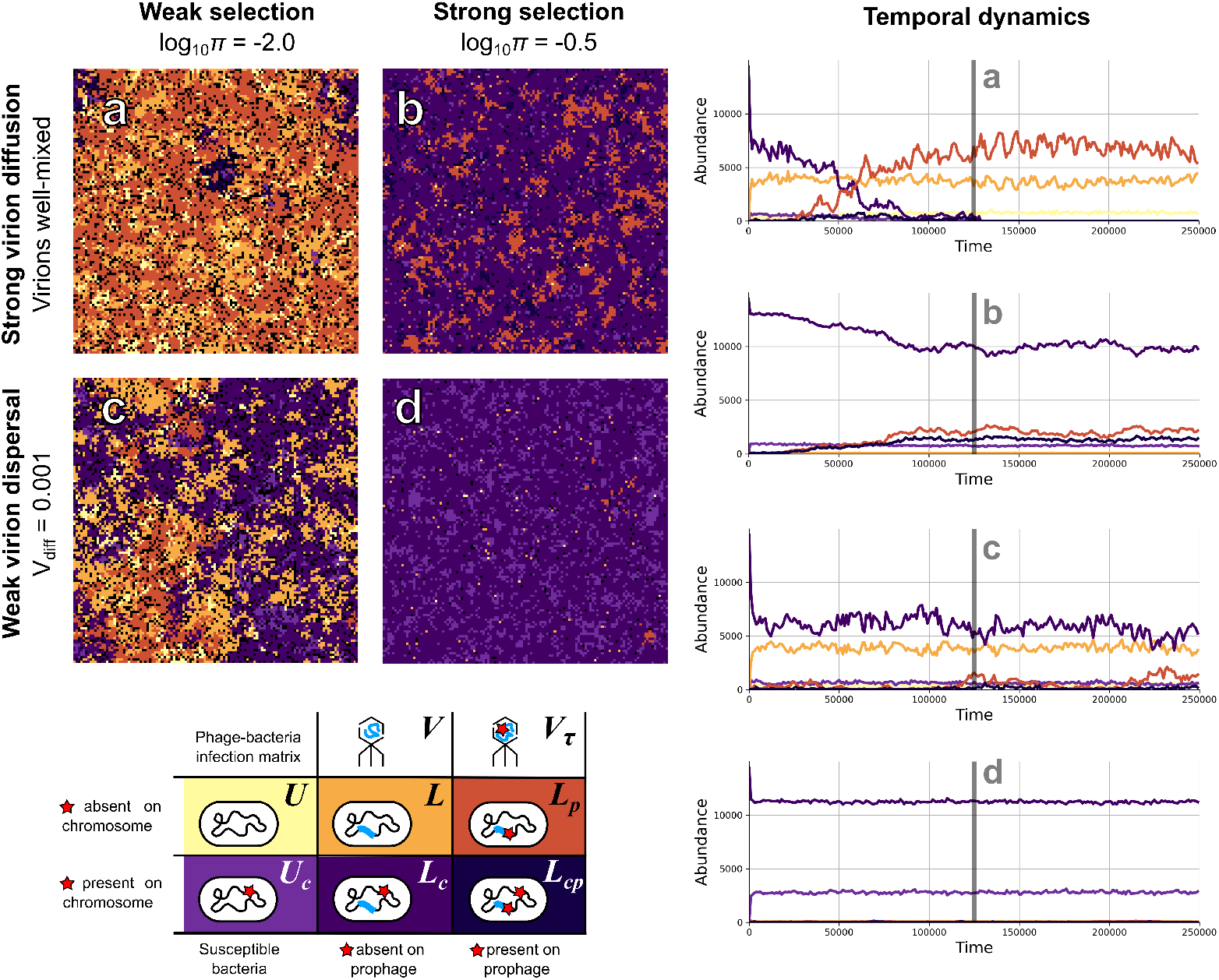
Snapshots of the spatial model at different time-points of a single run of the simulation. The four cases shown here give a side-by-side comparison to illustrate the limitations of lowering virion dispersal and increasing selection coefficient in the successful invasion of the phage-encoded gene. These simulations were run for a more extreme case with a higher gene loss rate (*µ* = 5 *×* 10^−4^) and lower virion decay (*µ* = 5 *×* 10^−3^) to clearly illustrate the spatial patterns.

While both chromosomal and phage-encoded versions of the gene can invade these clusters, the phage locus has a unique advantage: virion dispersal allows it to reach distant, gene-free regions that the chromosomal gene cannot access as easily. This effect is evident under weak selection and high virion diffusion (Figure 5(a)), where phage-mediated transmission facilitates successful invasion. However, when virion diffusion is limited (Figure 5 (c,d)), the advantage of the phage locus is lost and virions remain localised, unsuccessful in reaching susceptible cells in distant gene-free patches.

In contrast, strong selection (Figure 5(b)) eliminates gene-free regions through selective sweeps. Under these conditions, the chromosomal locus dominates due to its mutational robustness. Interestingly, under strong selection with high virion diffusion (Figure 5(b)), the system stabilises into a persistent coexistence of the chromosomal and phage-encoded gene, as neither is able to fully outcompete the other.

In summary, the success of the phage-encoded gene depends critically on two factors: effective virion dispersal and the continued presence of gene-free populations. These conditions allow the phage-encoded gene to utilise spatial heterogeneity and outcompete the chromosomal gene.

Based on these findings, we hypothesise the following.

*Accessory genes in bacteria can evolve to be associated with phages because of its ability to disperse these genes to favourable niches through virion particles*.

See Figure 6 for an illustration of the hypothesis.

**Figure 6.**
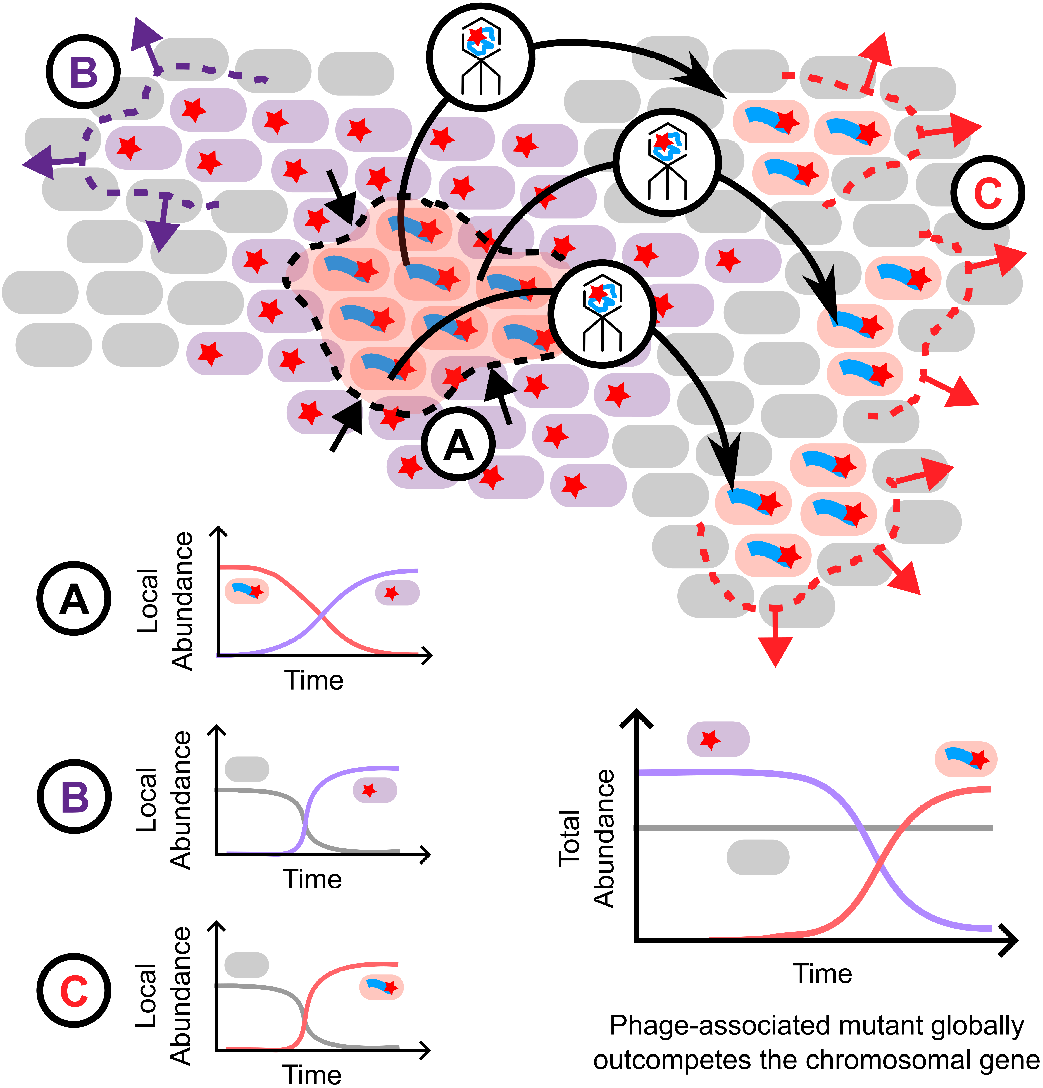
Dispersal hypothesis illustrated for phage-encoded beneficial traits. When a chromosomal gene can locally outcompete its phage-associated counterpart, but both types can locally outcompete noncarriers, the phage-associated gene gains a unique dispersal advantage. By diffusing through virion particles, it increases the mixing of carrier and non-carrier genotypes and thereby has a higher effective rate of selection over the chromosomal gene. This enables the phage-associated gene to outcompete the chromosomal gene on a larger spatial scale.

The occurrence of these favourable niches can be attributed to the weak selection pressure on the gene in a spatially structured cell population. This mechanism sets apart phage-encoded genes from genes associated with more locally transmitted MGEs such as plasmids and could potentially explain why phages differ from other MGEs in terms of the composition of their cargo.

## Discussion

Our study presents a suite of theoretical models to investigate the evolution of phage-associated genes originating from chromosomal loci. We verify that chromosomal mutational robustness and competition with non-carrier phages create significant barriers to the evolution of phage-encoded accessory genes. However, we find that spatial structure – both between and within populations – can help overcome these barriers. We propose that environmental heterogeneity can create selection pressures driving phage-gene associations, while spatial structure within bacterial populations can promote association via virion-mediated gene dispersal, without any regulatory coupling. In other words, we identified exogenous factors that can explain the observed association between prophages and virulence genes in contrast to previously hypothesised endogenous mechanisms.

### Co-regulation of virulence and prophage induction

Our findings suggest that when prophage induction is elevated in specific host environments, genes conferring local fitness benefits readily become associated with these prophages. This association is facilitated by the continual influx of susceptible, non-carrier cells from external environments. Previous work has shown that the regulation of phage-encoded virulence genes is highly complex and context-dependent. Although certain virulence genes may appear to be regulated independently of prophages under controlled in vitro conditions, bacterial cells in nature are exposed to diverse mixtures of signalling factors that vary widely across environments. Migration between such environments may therefore promote linkage between prophages and virulence genes through emergent correlations in their expression. Notably, environmental conditions characteristic of infection sites such as the gut (Henrot and Petit 2022) and urinary tract (Miller-Ensminger et al. 2020) are known to increase rates of prophage induction. In light of our results, it would be valuable to experimentally test whether prophages induced in these contexts exhibit a higher tendency to carry virulence genes.

In our analysis, we assumed symmetric cell densities and migration rates between the two environments, but in reality, these factors may vary. Exploring how different metapopulation structures influence this evolutionary process would be an interesting direction for future research. Nonetheless, our findings help explain the empirical tendency of phages to encode virulence factors (Boyd and Brüssow 2002; Abedon and LeJeune 2007; Schroven, Aertsen, and Lavigne 2021; Takeuchi, Hamada-Zhu, and Suzuki 2023). More broadly, our model applies to any weakly selected beneficial trait. Virulence genes fit this criterion due to their known tendency to share the benefits of pathogenicity with surrounding bacteria (Nogueira et al. 2009). The requirement for weak selection also aligns with a scenario where increasing host immunity decreases the selective pressure on virulence genes.

### Prophages are distinct from other MGEs as vehicles of HGT

Phages offer distinct advantages over plasmids in associating with virulence genes due to their lifecycle. Unlike plasmids, which rely on contact-dependent conjugation, phages achieve horizontal transfer through virion dispersal, and induction allows phages to synchronise gene transfer with adaptive stress responses. While cross-species transfer has been proposed to facilitate plasmid-associated gene evolution (Lehtinen, Huisman, and Bonhoeffer 2021), temperate phages may be more host-specific due to receptor-mediated infection and site-specific integration (Moura de Sousa et al. 2021; López-Leal et al. 2022). While plasmids carrying antimicrobial resistance genes may have host ranges spanning different genera or beyond (Coluzzi and Eduardo PC Rocha 2025), broad host range temperate phages are typically limited to strains within the same species or genus (Ford et al. 2021; Hammerl et al. 2022; Ilyas et al. 2022). Furthermore, evidence suggests that phage-associated virulence genes originate from chromosomal loci via erroneous excision of prophages (Brüssow, Canchaya, and Hardt 2004). While priority effect has been shown to drive plasmid-gene associations (Lehtinen, Huisman, and Bonhoeffer 2021), our models suggest that localised priority effects in spatially structured populations may instead facilitate the evolution of phage-encoded genes.

### Evolutionary timescales

Weak selection emerges as a key condition for the evolution of MGE-borne genes (Dijk and Hogeweg 2016; van Dijk et al. 2020; Lehtinen, Huisman, and Bonhoeffer 2021). Strong selection, typically associated with essential genes, instead favours chromosomal association due to greater stability and reliability. Under weak selection, frequent gene loss allows non-carrier and carrier lineages to coexist, making mobility advantageous by enabling gene spread independently of host reproduction. Due to weak selection, our simulations show that fixation of mobile genes occurs over long timescales, amidst significant demographic noise (see Supplementary Figures 1 and 2). Stable persistence, or sometimes fixation, of prophage-borne genes arises across a broad range of selection strengths when virion dispersal is high. However, when dispersal is limited, selection must be weak relative to gene loss, since the benefit of dispersal depends on discovering non-carrier regions; thus, higher abundances of non-carriers are required in the spatial model. To demonstrate this mechanism, we used gene-loss rates that may exceed typical chromosomal values, which vary widely across environments and genomic regions. This was a modelling choice to keep the framework simple and computation time tractable. Nevertheless, the dispersal mechanism is generalisable: it can operate under lower loss rates if selection is proportionally weaker, and also under stronger selection if coexistence of non-carriers is maintained by other processes. For example, cell influx, which may occur more frequently than gene loss, can also help non-carriers persist. Finally, the effectiveness of dispersal also depends on virion decay. Under favourable conditions, in the absence of UV-radiation for instance, virions can persist much longer than assumed here and may therefore disperse across greater distances.

### Spatial privatisation of benefits

In a spatially structured population, cooperation can persist through spatial privatisation – benefits being shared primarily within local neighbourhoods. When we restrict sharing of benefits from carrier cells to a radial neighbourhood, we found that phage-gene association evolved more easily with increasing virion dispersal as well as increasing the range of cooperation (see Supplementary Figure 3). This is because increasing the radius of sharing benefits increases the chance of non-carrier cells receiving the benefit and lowers the selection coefficient acting on the gene. Therefore, we highlight the generalisability of our dispersal hypothesis to any weakly selected gene in a spatially structured bacterial population.

For illustrative purposes, our models rely on several simplifying assumptions, deviating from the realities of natural systems. One notable assumption is superinfection immunity. In natural bacterial populations, particularly among pathogens, multiple prophages often coexist within a single host cell (Brüssow, Canchaya, and Hardt 2004; López-Leal et al. 2022). These prophages can undergo high order interactions and genetic exchange that reflect in dynamics at higher levels of biological systems (Moura de Sousa et al. 2021; Wendling 2023). While embedding these interactions in ODE models inflates the number of equations, it will be interesting to relax superinfection immunity in individual-based models in the future. These models may also include other processes we have thus far neglected, such as retention of partial immunity after phage loss (Arias et al. 2022). While these unmodelled complexities are potentially significant, they do not invalidate the core insights gained from our current model.

A further limitation of our model is that a chromosomal gene is indistinguishable from a gene carried by an inactivated prophage, i.e. a prophage lacking functional lysis genes and therefore unable to be induced. Consequently, we assumed that prophage loss mutations always entail the loss of any associated accessory gene. While this is reasonable in the case of prophage deletions, it does not necessarily hold for prophage inactivation, where accessory gene function can be retained. To address this, we also simulated a scenario in which the accessory gene is preserved as a chromosomal locus following prophage loss (see Supplementary Figures 4 and 5). With this modification, prophage–gene linkage evolved but persisted in small abundances in both the two-compartment model as well as the grid model. It remains unclear whether the abundant ‘chromosomal gene’ descended from an inactivated prophage–borne gene due to dispersal benefits from its inducible form. Further work will be needed to test this possibility.

In summary, our findings highlight spatial structure as a critical mechanism for the evolution of phage-encoded accessory genes. By explicitly modelling phage-specific dynamics, rather than MGEs in the abstract, we provide a framework to understand why phages are more likely to associate with virulence genes (Takeuchi, Hamada-Zhu, and Suzuki 2023). Our work provides mechanistic insights into the unique evolutionary role of temperate phages in bacterial adaptability, and their role in the persistence of virulence.

## Authors’ contributions

S.A.N. conceived the model with input from B.D. and designed and programmed the model, ran simulations, generated figures from the output data and drafted the initial manuscript. All authors gave final approval for publication and agree to be held accountable for the work performed therein.

## Acknowledgements

We thank Paulien Hogeweg, Otto Cordero, Nobuto Takeuchi and Hilje Doekes for valuable discussions and feedback on earlier versions of this work.

## Software

Simulations of the analytical model were run using Jupyter notebook (Kluyver et al. 2016), and the packages Matplotlib (Hunter 2007), NumPy (Harris et al. 2020), SciPy (Virtanen et al. 2020), and Pandas (McKinney 2010). The spatial model is implemented using Cacatoo (Dijk 2022), and an interactive version is available at https://sultannazir.github.io/bacteria_prophage_codes/.

## Data availability

All code and data for the model are available online at https://github.com/sultannazir/bacteria_prophage_codes.

