## Supplementary material for "Viral lifecycle dynamics and spatial structure explain why accessory genes are associated with temperate phages"

October 14, 2025

#### A Ordinary Differential Equations

The described model can be expressed in the form of differential equations. We use the notations used for the cell genotypes as non-negative real-valued variables to measure their corresponding abundances in the population. Therefore, the total cell population size ($N$) and the fractional abundance of carriers ($f_\tau$) can be expressed as

$$N = U + U_c + L + L_c + L_p + L_{cp}$$
$$f_\tau = (U_c + L_c + L_p + L_{cp})/N$$

Similarly, we use $V$ and $V_\tau$ to denote the abundances of non-carrier and carrier virion particles. Hence, we describe the temporal dynamics of the cell and phage abundances using the following system of differential equations:

$$
\begin{aligned}
\frac{dU}{dt} &= U\left(1 - d - N + (1-\pi)\phi f_\tau\right) && -\beta U(V + V_\tau) && +\lambda(L + L_p) + \mu U_c && \\
\frac{dU_c}{dt} &= U_c\left(1 - d - N + \pi\phi + (1-\pi)\phi f_\tau\right) && -\beta U_c(V + V_\tau) && +\lambda(L_c + L_{cp}) - \mu U_c && \\
\frac{dL}{dt} &= L\left(1 - d - N + (1-\pi)\phi f_\tau\right) && +\beta U V && -\lambda L + \mu(L_c + L_p) && -\alpha L \\
\frac{dL_c}{dt} &= L_c\left(1 - d - N + \pi\phi + (1-\pi)\phi f_\tau\right) && +\beta U_c V && -(\lambda+\mu)L_c + \mu L_{cp} && -\alpha L_c \\
\frac{dL_p}{dt} &= L_p\left(1 - d - N + \pi\phi + (1-\pi)\phi f_\tau\right) && +\beta U V_\tau && -(\lambda+\mu)L_p + \mu L_{cp} && -\alpha L_p \\
\frac{dL_{cp}}{dt} &= \underbrace{L_{cp}\left(1 - d - N + \pi\phi + (1-\pi)\phi f_\tau\right)}_{\text{Growth}} && +\underbrace{\beta U_c V_\tau}_{\text{Infection}} && -\underbrace{(\lambda+2\mu)L_{cp}}_{\text{Mutations}} && -\underbrace{\alpha L_{cp}}_{\text{Induction}}
\end{aligned}
$$

$$
\begin{aligned}
\frac{dV}{dt} &= r_V\alpha(L + L_c) && -\beta(U + U_c)V && -\delta V \\
\frac{dV_\tau}{dt} &= \underbrace{r_V\alpha(L_p + L_{cp})}_{\text{Lytic release}} && -\underbrace{\beta(U + U_c)V_\tau}_{\text{Infection}} && -\underbrace{\delta V_\tau}_{\text{Decay}}
\end{aligned}
$$

### B Model parameters

| **Variable** | **Meaning** | **Value (ODE/IBM)** |
|---|---|---|
| $U$ | Abundance of non-carrier susceptible cells | $[0, \infty)$ / $\{0, 1, 2, 3...\}$ |
| $U_c$ | Abundance of susceptible cells carrying gene on the chromosomal locus | $[0, \infty)$ / $\{0, 1, 2, 3...\}$ |
| $L$ | Abundance of non-carrier lysogens | $[0, \infty)$ / $\{0, 1, 2, 3...\}$ |
| $L_c$ | Abundance of lysogens carrying the gene on the chromosomal locus | $[0, \infty)$ / $\{0, 1, 2, 3...\}$ |
| $L_p$ | Abundance of lysogens carrying the gene on the phage locus | $[0, \infty)$ / $\{0, 1, 2, 3...\}$ |
| $L_{cp}$ | Abundance of lysogens carrying the gene on both loci | $[0, \infty)$ / $\{0, 1, 2, 3...\}$ |
| $V$ | Abundance of extracellular non-carrier virions | $[0, \infty)$ / $\{0, 1, 2, 3...\}$ |
| $V_\tau$ | Abundance of extracellular carrier virions | $[0, \infty)$ / $\{0, 1, 2, 3...\}$ |
| **Parameter** | **Meaning** | **Value (ODE/IBM)** |
| $\phi$ | Ratio of maximum increase in carrying capacity to minimum carrying capacity | $1$ / $1$ |
| $b$ | Intrinsic birth rate | $1$ / $0.5$ |
| $d$ | Intrinsic death rate | $0.02$ / $5 \times 10^{-3}$ |
| $\pi$ | Fraction of benefits privatised by carriers of gene | $[10^{-4.5}, 10^{-1.0}]$ / $[10^{-3.5}, 10^{-2.0}]$ |
| $\beta$ | Rate of phage infection | $0.04$ / $0.01$ |
| $\alpha$ | Rate of prophage induction and lysis | $[10^{-4.5}, 10^{-1.0}]$ / $10^{-3}$ |
| $r_V$ | Burst size | $30$ / $30$ |
| $\delta$ | Rate of virion decay | $0.04$ / $0.01$ |
| $\mu$ | Rate of gene loss | $2 \times 10^{-4}$ / $5 \times 10^{-5}$ |
| $\lambda$ | Rate of phage loss | $2 \times 10^{-3}$ / $5 \times 10^{-4}$ |
| $m$ | Rate of migration | $0$ (one compt.), $10^{-3}$ (two compt.) / n.a. for IBM |
| $V_{\text{diff}}$ | Rate of virion diffusion | $\{10^{-3}, 10^{-1}, \infty\}$ (IBM) |

Table 1: Variables and parameters of the model, their meaning and the values used. The variables are non-dimensionalised in the ODEs and the rate parameters are scaled when switching between the ODE and IBM. The details of these scales are given in Appendix C

### C Well-mixed approximation of the spatial model

Let $x$ be the number of cells on the grid of a certain genotype and $k$ be the minimum carrying capacity, i.e. the grid size is $k(1+\phi)$. Then, the probability of finding an empty grid point is

$$\mathbb{P}(\text{empty grid point}) = 1 - \frac{n}{k(1+\phi)}$$

The average death rate of a genotype will be

$$\frac{\sum_{\text{genotype}} \mathbb{P}_{\text{death}}(\pi)}{\sum_{\text{genotype}} 1} = \begin{cases} d_s + \Delta d\Big(1 - \pi - (1-\pi) f_\tau\Big), & \text{genotype is a carrier} \\ d_s + \Delta d\Big(1 - (1-\pi) f_\tau\Big), & \text{genotype is not a carrier} \end{cases}$$

The the well-mixed growth dynamics of a carrier genotype is given by

$$\frac{dx}{dt'} = \text{Avg. \#births into empty grid points} - \text{Avg. \# deaths}$$
$$\frac{dx}{dt'} = bx\Big(1 - \frac{n}{k(1+\phi)}\Big) - x\Big(d + \Delta d\Big(1 - \pi - (1-\pi) f_\tau\Big)\Big)$$
$$\frac{dx}{dt'} = \frac{b}{1+\phi} x\Big(1 + \phi - \frac{n}{k} - d_s\frac{1+\phi}{b} - \Delta d\frac{1+\phi}{b}\Big(1 - \pi - (1-\pi) f_\tau\Big)\Big)$$

We apply the following non-dimensionalisation

$$X = \frac{x}{k}, \quad N = \frac{n}{k}, \quad t = t'\frac{b}{1+\phi}$$

and the following substitutions

$$d_s = d\frac{b}{1+\phi}, \quad \Delta d = \frac{b\phi}{1+\phi}$$

to obtain

$$\frac{dX}{dt} = X(1 - N + \pi\phi + (1-\pi)\phi f_\tau)$$

which is the same growth dynamics we implemented in the ODEs. The same derivation can be done for the growth dynamics of a non-carrier genotype. Notice that time is scaled by a factor of $\frac{b}{1+\phi}$ in the ODE, i.e. one time-step in the IBM is equal to $\frac{1+\phi}{b}$ (which is equal to 4 for our choice of parameters) many time-steps of the ODE. Therefore, all the rate parameters, just like the intrinsic death rate, need to be adjusted to the time scale by this factor when switching between the models.

### D Selection- and transmission-mutation thresholds

To derive the persistence condition of the gene in the face of mutation, we can consider a simpler version of the model consisting only of two kinds of cells: carrier ($U$) and non-carrier ($U_c$).

$$\frac{dU}{dt} = U\left(1 - U - U_c + (1-\pi)\phi\frac{U_c}{U+U_c}\right) + \mu U_c$$
$$\frac{dU_c}{dt} = U_c\left(1 - U - U_c + \pi\phi + (1-\pi)\phi\frac{U_c}{U+U_c}\right) - \mu U_c$$

We obtain the condition that for a positive abundance of the carrier type at steady-state ($U_c(t=\infty) > 0$) requires the rate of selection to exceed the rate of mutation:

$$\pi\phi > \mu$$

*Proof.* Consider the non-dimensionalised version of the model such that $u = U/N$ and $u_c = U_c/N$, where $N = U + U_c$.

The equations then become

$$\frac{du}{dt} = -\pi\phi u u_c + \mu u_c$$
$$\frac{du_c}{dt} = \pi\phi u u_c - \mu u_c$$

Solving for steady state, the non-trivial solutions are $u^\infty = \frac{\mu}{\pi\phi}, u_c^\infty = 1 - \frac{\mu}{\pi\phi}$, and $u^\infty = 1, u_c^\infty = 0$.

We can infer that a stable positive solution for $u_c^\infty$ exists only when $\pi\phi > \mu$. □

Now to derive the persistence condition of the MGE, let us look at a simpler version of the model consisting only of two kinds of bacteria: uninfected ($U$) and infected ($L$). First, let us consider a plasmid.

$$\frac{dU}{dt} = U(1 - U - L) - \beta UL + \lambda L$$
$$\frac{dL}{dt} = L(1 - c - U - L) + \beta UL - \lambda L$$

We obtain the condition that the plasmid will persist in the population ($L(t=\infty) > 0$) if the conjugation rate is higher than the sum of rate of segregational loss and cost of carrying the plasmid:

$$\beta > \lambda + c$$

*Proof.* The plasmid will persist if it has a positive growth rate when it is rare in the population, i.e. the susceptible cells are maximally abundant ($U = 1$) and the infected cell abundance is close to zero ($L^2 \simeq 0$).

Then,

$$\frac{dL}{dt} > 0 \implies \beta - \lambda - c > 0$$

$\square$

When the MGE is a phage, we will consider an equation describing the virion abundance $(V)$ as well.

$$\begin{aligned}
\frac{dU}{dt} &= U(1 - U - L) - \beta UV + \lambda L \\
\frac{dL}{dt} &= L(1 - U - L) + \beta UV - \lambda L - \alpha L \\
\frac{dV}{dt} &= r_V \alpha L - \beta UV - \delta V
\end{aligned}$$

We obtain the condition that the phage will persist in the population $(L(t = \infty) > 0)$ if the lysis rate is above a certain threshold:

$$\alpha > \frac{\lambda(\beta + \delta)}{\beta(r_V - 1) - \delta}$$

*Proof.* By explicitly considering the extracellular virions, we have modelled an infection cycle with two stages: the prophage stage and the virion stage. For a two-stage infection cycle, the net basic reproductive number of the infection, $R_0$ is given by the product of $R_0$ of each step. Therefore, the phage will persist if the net $R_0$ is greater than 1.

$$\begin{aligned}
R_0 &= R_0(\text{prophage}) \times R_0(\text{virion}) \\
&= \frac{\text{max. infection rate}}{\text{phage loss rate} + \text{lysis rate}} \times \frac{\text{burst size} \times \text{lysis rate}}{\text{max. infection rate} + \text{decay rate}} \\
&= \frac{r_V \alpha \beta}{(\lambda + \alpha)(\beta + \delta)}
\end{aligned}$$

Therefore,

$$R_0 > 1 \implies \alpha > \frac{\lambda(\beta + \delta)}{\beta(r_V - 1) - \delta}$$

$\square$

### E Modified ODEs

Let $m$ be the rate of migration between the 'host' and the 'outside world' compartments. Then the ODEs describing the dynamics in the outside world (variables denoted with a superscript $o$ represent abundances in the outside world).

$$
\begin{aligned}
\frac{dU^o}{dt} &= U^o(1+\phi-d-N^o) - \beta U^o(V^o+V_\tau^o) + \lambda(L^o+L_p^o) + \mu U_c^o + m(U-U^o)\\
\frac{dU_c^o}{dt} &= U_c^o(1+\phi-d-N^o) - \beta U_c^o(V^o+V_\tau^o) + \lambda(L_c^o+L_{cp}^o) - \mu U_c^o + m(U_c-U_c^o)\\
\frac{dL^o}{dt} &= L^o(1+\phi-d-N^o) + \beta U^o V^o - (\alpha_{\text{out}}+\lambda)L^o + \mu(L_c^o+L_p^o) + m(L-L^o)\\
\frac{dL_c^o}{dt} &= L_c^o(1+\phi-d-N^o) + \beta U_c^o V^o - (\alpha_{\text{out}}+\lambda+\mu)L_c^o + \mu L_{cp}^o + m(L_c-L_c^o)\\
\frac{dL_p^o}{dt} &= L_p^o(1+\phi-d-N^o) + \beta U^o V_\tau^o - (\alpha_{\text{out}}+\lambda+\mu)L_p^o + \mu L_{cp}^o + m(L_p-L_p^o)\\
\frac{dL_{cp}^o}{dt} &= L_{cp}^o(1+\phi-d-N^o) + \beta U_c^o V_\tau^o - (\alpha_{\text{out}}+\lambda+2\mu)L_p^o + m(L_{cp}-L_{cp}^o)\\
\frac{dV^o}{dt} &= r_V\alpha_{\text{out}}(L^o+L_c^o) - \beta(U^o+U_c^o)V^o - \delta V^o + m(V-V^o)\\
\frac{dV_\tau^o}{dt} &= r_V\alpha_{\text{out}}(L_p^o+L_{cp}^o) - \beta(U^o+U_c^o)V_\tau^o - \delta V_\tau^o + m(V_\tau-V_\tau^o)
\end{aligned}
$$

where $N^o = U^o + U_c^o + L^o + L_c^o + L_p^o + L_{cp}^o$. The only modification to the system of ODEs for the host compartment is the addition of a migration term $+m(X^o - X)$ to the ODE for the corresponding $X$ genotype/virion abundance variable.

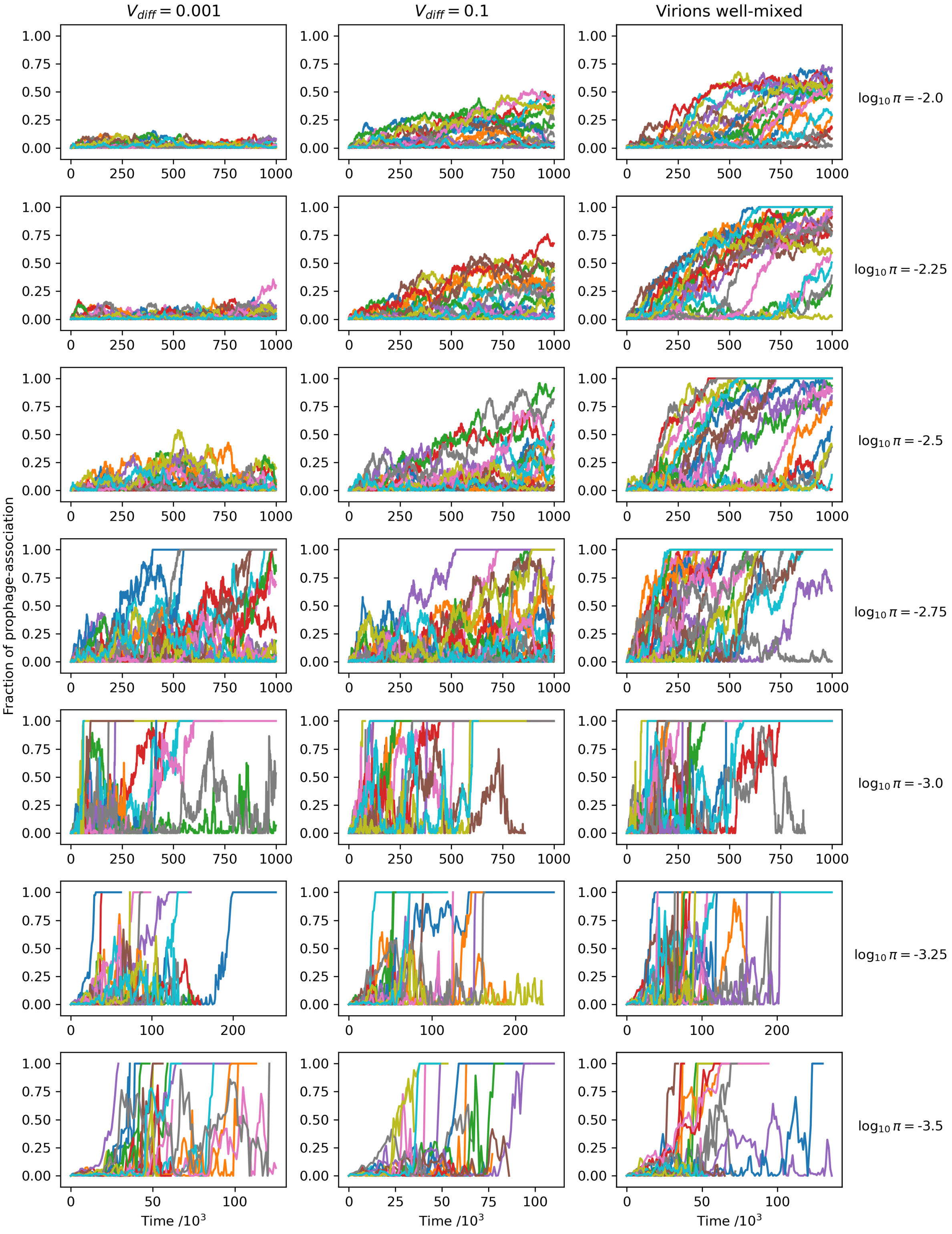


Supplementary Figure. 1: Cells are stationary on the grid: Time-series plots of the fraction of phage-associated genes for 20 replicates per value of privatisation (rows) and virion diffusion rate (columns).

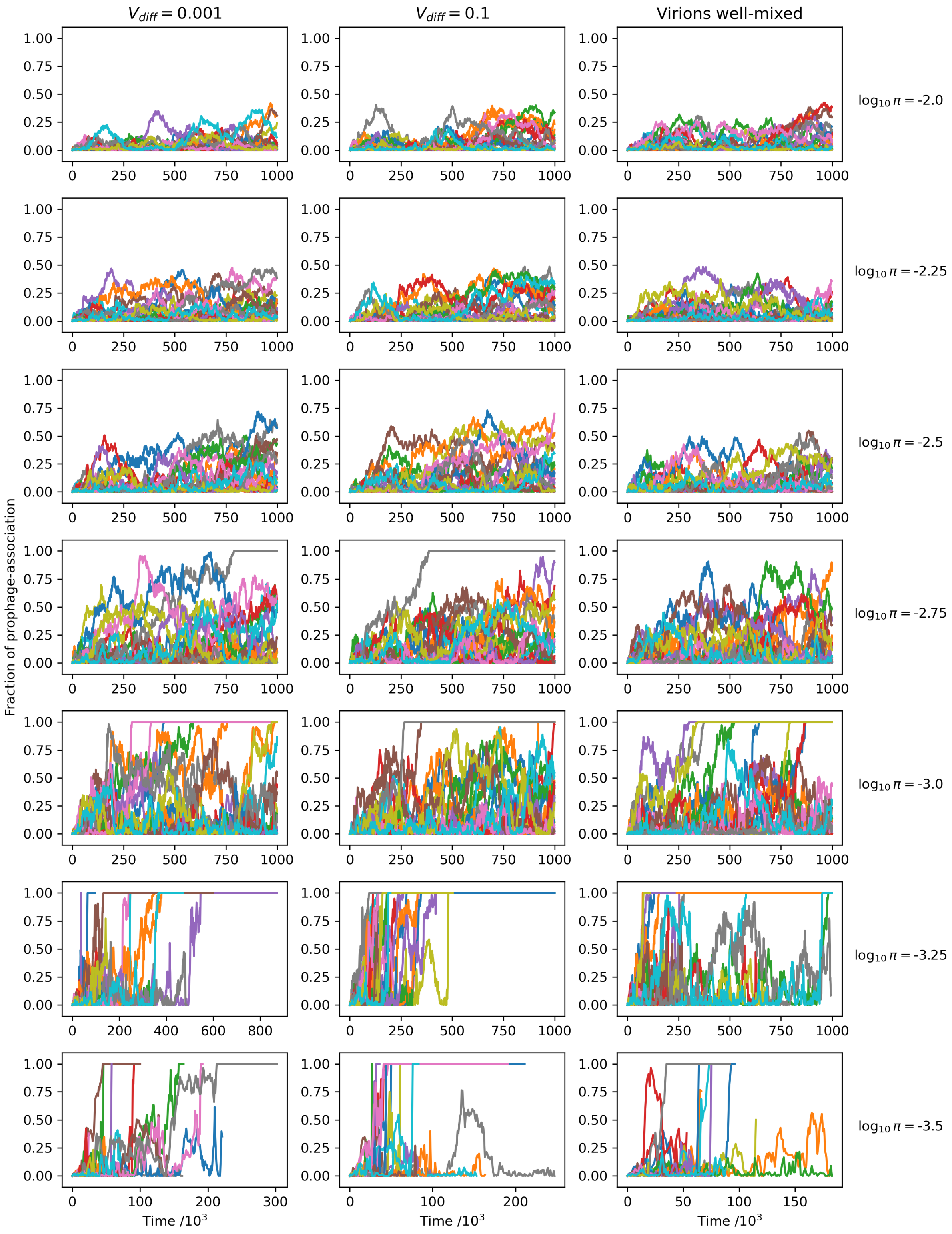


Supplementary Figure. 2: Cells are well-mixed on the grid: Time-series plots of the fraction of phage-associated genes for 20 replicates per value of privatisation (rows) and virion diffusion rate (columns).

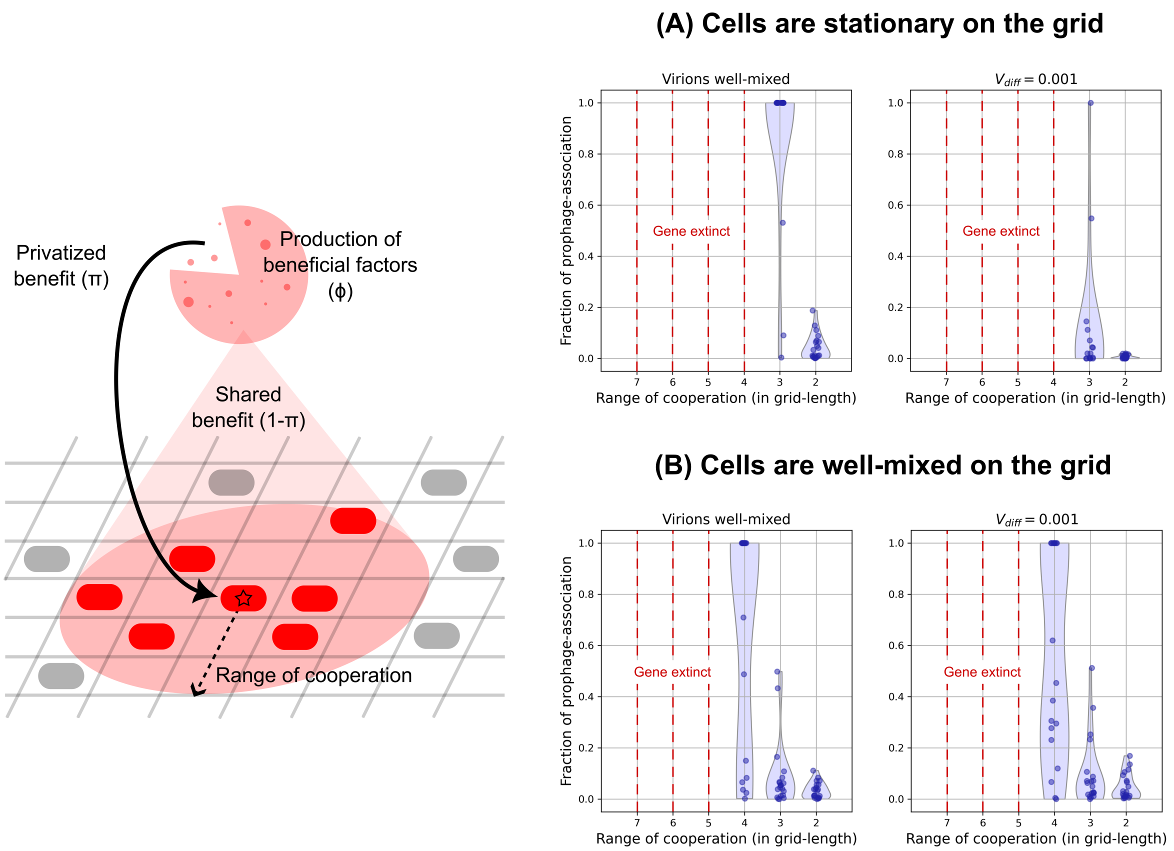


Supplementary Figure. 3: (Left) In a spatial model, privatization can be modelled mechanistically through spatial privatization, i.e. the benefits of expression of a cooperative trait is only experienced by cells in an immediate neighbourhood. Therefore, we introduce a parameter "Range of cooperation" which measures the maximum distance of a cell from a carrier cell in order to obtain benefits from the latter. Therefore, decreasing this radius increases the privatization. Furthermore, since a cell is likely surrounded by similar genotype, decreasing the radius also increases the selection pressure acting on the gene. (Right) With the mechanistic implementation of privatization, we yet again observe that the chance of prophage-association increases with increasing virion dispersal and decreasing selection coefficient (increasing range of cooperation). In comparison with the case where the cells are well-mixed, we can clearly see that spatial structure is required for this trend. We also see that gene extinction happens sooner with spatial structure and that the effect is stochasticity is much stronger. This is because at the local level, the carrier genotypes have the same growth rate as the non-carrier (cheater) genotypes and hence evolves towards gene extinction. Selection in this model occurs at a higher (group-) level. This sufficiently reduces the effective population size on which selection is acting - leading to increased genetic drift. To increase the effective population size, we lowered the parameter $\phi$ to $0.1$ and adjusted other parameter values accordingly. These figures were generated for different values of certain parameters: $\mu = 5 \times 10^{-4}$, $\alpha = 5 \times 10^{-4}$ and $\delta = 5 \times 10^{-3}$

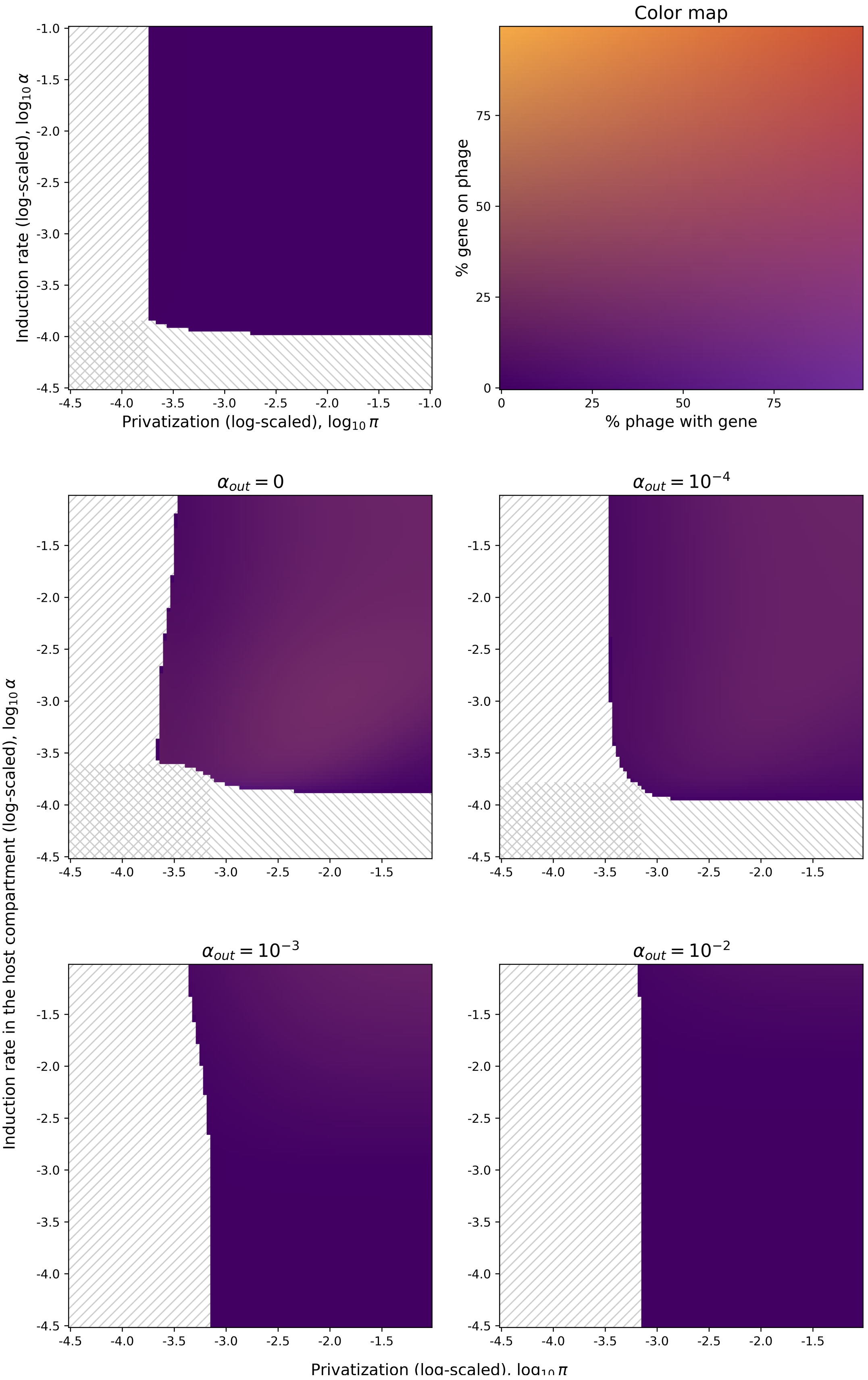


Supplementary Figure. 4: ODE simulation: prophage inactivation with gene retention. $L_p$ genotype becomes $U_c$ upon prophage loss. Plots show final degree of gene-phage association in a parameter-sweep across privatisation and induction rate.

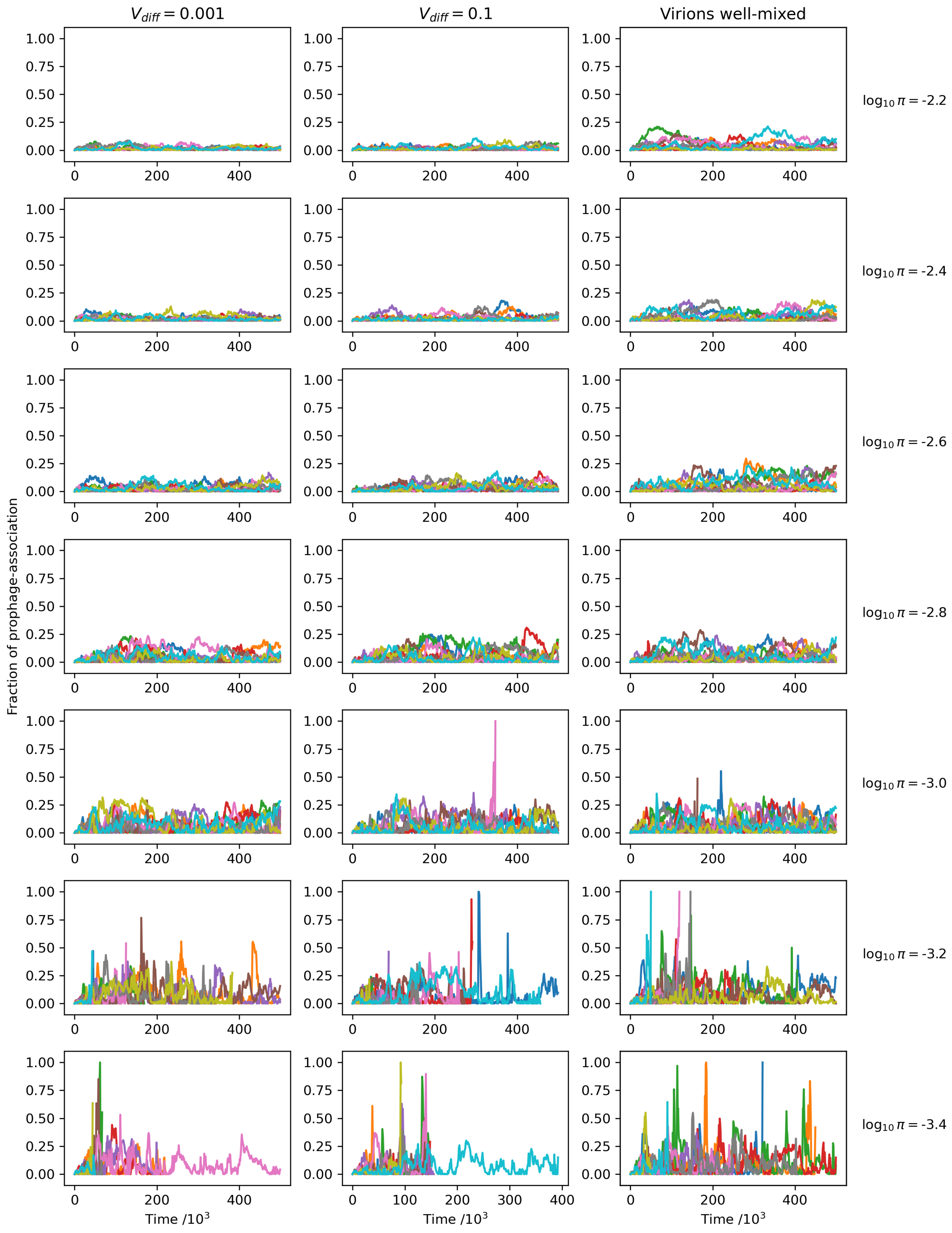


Supplementary Figure. 5: Grid simulation: prophage inactivation with gene retention. $L_p$ genotype becomes $U_c$ upon prophage loss. Plots show time-series of the fraction of phage-associated genes for 20 replicates per value of privatisation (rows) and virion diffusion rate (columns).
